# Development of a first-in-class unimolecular dual GIP/GLP-2 analogue, GL-0001, for the treatment of bone fragility

**DOI:** 10.1101/2022.07.21.500659

**Authors:** Benoit Gobron, Malory Couchot, Nigel Irwin, Erick Legrand, Béatrice Bouvard, Guillaume Mabilleau

**Author notes:** **Correspondence to :** Dr Guillaume Mabilleau, Inserm UMR_S 1229 RMeS – REGOS Team, University of Angers, Institut de Biologie en santé, 4 rue Larrey, 49933 Angers, France.

## Abstract

Due to ageing of the population, bone frailty is dramatically increasing worldwide. Although some therapeutic options exist, they do not fully protect or prevent against the occurrence of new fractures. All current drugs approved for the treatment of bone fragility target bone mass. However, bone resistance to fracture is not solely due to bone mass but relies also on bone ECM material properties, i.e. the quality of the bone matrix component. Here, we introduce the first-in-class unimolecular dual GIP/GLP-2 analogues, GL-0001, that activate simultaneously the glucose-dependent insulinotropic polypeptide receptor (GIPr) and the glucagon-like peptide-2 receptor (GLP-2r). GL-0001 acts synergistically through a cAMP-LOX pathway to enhance collagen maturity. Furthermore, in mice with ovariectomy-induced bone fragility, GL-0001 prevented excess trabecular bone degradation at the appendicular skeleton and also enhanced bone ECM material properties through reduction of the degree of mineralization and augmentation in enzymatic collagen crosslinking. These results demonstrate that targeting bone ECM material properties is a viable option to enhance bone strength and opens an innovative pathway for the treatment of patients suffering of bone fragility.

## 1. INTRODUCTION

Due to ageing of the population, the occurrence of bone fragility and fracture has risen significantly worldwide and will continue in the future ^(1)^. Bone fragility develops as a concomitant decline in bone mass and bone quality. Bone mass is regulated by a complex balance between catabolic removal of quanta of bone material by osteoclasts and anabolic deposition of quanta of bone material by osteoblasts. All approved drug for the treatment of osteoporosis and bone fragility are based on either limiting osteoclast-mediated bone resorption or enhancing osteoblast-mediated bone formation. This is due to the paradigm that increasing bone mineral density, the clinical surrogate for bone mass assessment, will result in higher bone strength. Although this is partly true in osteoporosis, and despite all approved anti-osteoporotic drugs increase bone mineral density, the risk of bone fracture is only reduced by 30-40% in hip and long bones in treated individuals ^(2-5)^. This suggests that factors beyond bone mass are primordial for an optimum bone strength.

Bone quality is also compromised in bone fragility ^(6)^. Bone quality encompasses an ensemble of factors including bone microstructure, bone ECM material properties and resistance to crack propagation ^(7)^. Indeed, despite the relatively simple structure of the collagen triple helix, the biosynthesis of collagen molecules by osteoblasts undergoes multiple post-translational steps to ensure proper folding and resistance to deformation. Among all, enzymatic collagen crosslinking, ensured by lysyl hydroxylase and lysyl oxidase, has attracted attention ^(8)^. To date, only a few signaling pathways known to regulate bone ECM material properties, have been identified ^(9-12)^. Unravelling the mechanisms controlling bone ECM material properties should lead to the development of bone quality-specific drugs that could represent an interesting alternative to the existing medications.

The idea of selectively targeting bone ECM material properties to improve bone strength, rather than directly addressing low bone mass, has emerged as a result of pioneering works with two endogenous gut hormones, named GIP and GLP-2. Indeed, these peptides maintain optimal bone ECM material properties, through a cAMP-dependent pathway ^(13-16)^. This concept was also reinforced by the use of first generation GIPr- and GLP-2r-specific analogues in several mice models of bone fragility ^(17-21)^. Furthermore, head-to-head comparison in preclinical model of bone fragility suggested that single GIP or GLP-2 analogues improved resistance to fracture to the same magnitude level than observed with bisphosphonates ^(17,22)^. However, this effect was not observed with the closely related glucagon or glucagon-like peptide-1 peptides, highlighting the selective bone response to GIP and GLP-2 single analogues. This is particularly interesting with respect to the recent evidence that functional GIPr and GLP-2r are also present in human bone cells ^(23,24)^.

Based on these outcomes, we hypothesized that co-administration of single GIP or GLP-2 analogues would result in synergistic effects as compared with single parent molecules and would surpass the beneficial effects of bisphosphonates on bone resistance to fracture. This idea is reinforced by the recent report that GIP and GLP-2 exerts separate beneficial effects on bone turnover ^(25)^. Here, to overcome possible discrepancies in pharmacokinetic profiles of GIP and GLP-2 analogues we developed a series of unimolecular dual GIP/GLP-2 analogues and tested their capacity to bind and activate the human GIPr and GLP-2r, whilst also evaluating their capacity to enhance bone ECM material properties. Finally, we identified our lead molecule, a first in class unimolecular dual GIP/GLP-2 analogues, GL-0001. The latter dually binds the GIPr and GLP-2r, activates intracellular production of cAMP resulting in higher expression of lysyl oxidase and ultimately higher collagen cross-linking. GL-0001-mediated changes in bone ECM material properties also improved bone strength in a mouse model of ovariectomy-induced bone fragility. Unimolecular dual GIP/GLP-2 analogue represent an innovative and novel pathway to treat bone fragility that should be further explored for the treatment of bone fragility disorder.

## 2. MATERIAL AND METHODS

A table of key resources is supplied as supplementary table 1.

### 2.1. Osteoblast cultures

Murine MC3T3-E1 cells were grown in αMEM supplemented with 10% FBS, 100 UI/mL penicillin, and 100 μg/mL streptomycin in a humidified atmosphere enriched with 5% CO_2_ at 37 °C. For differentiation studies, cells were plated at a density of 15,000 cells/cm^2^, grown to confluence and differentiated in medium supplemented with 50 μg/mL ascorbic acid and several concentrations of peptides. Fourteen days later, osteoblast cultures were either fixed in absolute ethanol, scrapped off the culture dish and transferred onto BaF2 windows for evaluation of collagen maturity or prepared for gene expression.

For mechanistic investigations, administration of 50 µM 2’,5’-DDA ^(15)^ and 0.25 mM bAPN ^(26)^ were added in the differentiation medium and replenished every second day. For silencing experiments, at day 7 of differentiation studies, osteoblasts were transfected with 50 nM of siRNA targeting murine GIPr (assay ID s233873), murine GLP-2r (assay ID s211996), murine lysyl oxidase (assay ID s69290) or a control scrambled siRNA (ref 4390843) using lipofectamine RNAimax. Experiments were terminated at day 14 and ECM properties were evaluated as stated below.

Normal human osteoblasts were plated at a density of 5,000 cells/cm² in Clonetics™ OGM™ osteoblast growth medium. At 80% confluence, 200 nM hydrocortisone-21-hemissucinate was added to the culture to induce differentiation. Ninety-five days later, osteoblast cultures were fixed and processed as described above for murine osteoblasts.

### 2.2. Intracellular cAMP determination in osteoblast cultures

MC3T3-E1 cells were plated at a density of 15,000 cells/cm^2^ and grown for 48 h in αMEM supplemented with 10% FBS, 100 UI/mL penicillin, and 100 μg/mL streptomycin in a humidified atmosphere enriched with 5% CO_2_ at 37°C. After 48 h, cells were starved in αMEM supplemented with 0.5% BSA for 16 h and then incubated for 15 min in αMEM supplemented with 0.5% BSA and 1 mM IBMX prior to stimulation with peptides. After 45 min, cells were washed with ice-cold PBS, incubated in RIPA buffer, and centrifuged at 13,000 *g* for 10 min. Supernatants were collected and stored at − 80°C until use. Cyclic AMP determination was performed with a commercially available EIA kit according to the manufacturer’s recommendations.

### 2.3. Binding assay and receptor activation

Human GIPr and human GLP-2r cDNAs were cloned in pcDNA3.1 between XhoI and AgeI of the multiple cloning site. For binding assay, CHO-K1 cells were transfected with pcDNA3.1(hGIPr) or pcDNA3.1(hGLP-2r) using Lipofectamine™ 3000. Forty-eight hours later, various concentrations of analogues were added in the presence of either 10^−7^ M Fam-[D-Ala²]-GIP_1-30_ or 10^−6^ M Fam-[Gly^2^]-GLP-2 in αMEM supplemented with 0.1% BSA. These concentrations of fluorescent peptides were determined on preliminary assays and represented 100% binding. Equilibrium binding was achieved overnight at 4°C. Cells were then washed twice with assay buffer and solubilized in 0.1 M NaOH. Fluorescence was read with a SpectraMax M2 microplate reader with excitation wavelength set up at 490 nm and emission wavelength set up at 525 nm. Binding at the human GIP receptor or human GLP-2 receptor was achieved by non-linear regression analyses.

For activation assay, plasmids encoding the human GIPr, human GLP-2r and Epac-S-H74 probe, a validated cAMP FRET biosensor ^(27)^, were transfected with Lipofectamine™ 3000 into CHO-K1 cells. Forty-eight hours later, transfected cells were incubated in HEPES buffered saline in the presence of various concentrations of peptides for 30 minutes. Donor excitation was made at 460 nm, donor emission was collected at 480 nm and acceptor emission at 560 nm with a M2 microplate reader. FRET was expressed as the ratio between donor and acceptor signals. The FRET ratio was standardized at 1 with vehicle. An increase in FRET ratio suggests an augmentation in intracellular cAMP levels.

### 2.4. Gene expression

For osteoblast cultures, total RNA was extracted after rinsing cultures with PBS. Ex vivo gene expression analysis was performed after crushing left tibia in a liquid nitrogen-cooled biopulverizer. Nucleozol (Macherey-Nagel, Hoerdt, France) was added on top of the cell layer/bone powder and total RNA were purified with Nucleospin RNA set nucleozol column (Macherey-Nagel) according to the manufacturer recommendations. Total RNA was reversed transcribed using Maxima first strand cDNA synthesis kit. Real-time qPCR was performed using TaqMan™ Fast advanced master mix and TaqMan Gene Expression Assays for *Lox* (Mm00495386_m1), *Col1a1* (Mm00801666_g1), *Alpl* (Mm00475834_m1) and *Plod2* (Mm00478767_m1). The *B2m* endogenous control (Mm00437762_m1) was used for normalization using the 2^-ΔΔCT^ method.

### 2.5. Osteoclast cultures

Murine Raw 264.7 cells were plated at a concentration of 125,000 cells/cm² in osteoclast medium containing αMEM supplemented with 10% FBS, 2 mM L-glutamine, 100 U/ml penicillin and 100 µg/ml streptomycin. To generate osteoclasts from Raw264.7 cells, osteoclast medium was enriched with 25 ng/ml sRANKL and several concentrations of peptides were then added to test their effects on osteoclast differentiation. After five days of culture, cytochemical TRAcP staining was performed to evidence the presence of osteoclast cells (TRAcP positive cells with more than 3 nuclei).

Isolated human PBMCs were plated at a concentration of 100,000 cells/cm² in osteoclast medium enriched with 25 ng/ml recombinant human M-CSF, 25 ng/ml recombinant human sRANKL (added at day 7) and various concentrations (10^−10^ – 10^−8^ M) of peptides (added at day 7). Cultures made for assessment of osteoclast resorption were plated on collagen-coated plastic plates. Cultures were terminated after 14 days to assess the extent of osteoclast formation (TRAcP positive cells with more than 3 nuclei) or at 21 days to assess the extent of osteoclast resorption (see below). All factors were replenished every 2-3 days.

The expression of TRAcP was examined cytochemically, as described previously ^(28)^ using naphtol AS-BI-phosphate as substrate and fast violet B as the diazonium salt. Cells were counterstained with 4, 6-diamidino-2-phenylindole (DAPI).

For assessment of osteoclast resorption, human osteoclasts cultured on collagen-coated plates were incubated with collagenase at day 20. The cell suspension was passed through a cell strainer with 40 µm pores, and the fraction retained in the strainer was plated on dentine slices at a concentration of 5,000 cells/cm² and cultured for an additional 24 hrs. After discarding osteoclasts, dentine slices were stained with 0.5% (v/v) toluidine blue (pH 5.0) prior to examination by light microscopy for the presence of lacunar resorption, the extent of surface erosion on dentine slices was determined using image analysis as described previously _(29)_.

### 2.6. Animals

All procedures were carried out in accordance with the European Union Directive 2010/63/EU for animal experiments and were approved by the regional ethical committee for animal use (authorization CEEA-PdL06-01740.01). Briefly, bilateral OVX was performed in 32 BALB/c (BALB/cJRj) mice at 12 weeks of age under general anesthesia. At 16 weeks of age, osmotic minipumps were implanted subcutaneously between the two scapulae in 24 OVX mice that were randomly allocated into either saline (OVX+saline, n=8), 25 nmoles/kg/day GL-0001 (OVX+GL-0001, n=8) or 25 nmoles/kg/day GL-0007 (OVX+GL-0007, n=8). This dose of unimolecular dual GIP/GLP-2 analogues was based on previous studies performed with single parent peptides ^(17,22)^. Eight ovariectomized BALB/c mice received an intravenous injection of zoledronic acid (100 µg/kg) and were implanted subcutaneously with an osmotic minipump filled with saline at 16 weeks of age (OVX+Zoledronic acid). Eight sham operated female BALB/c mice with the same age and implanted with osmotic minipumps delivering saline were used as negative controls (Sham+saline). Osmotic minipump were exchanged after 4 weeks. All animals received an intraperitoneal administration of calcein green (10 mg/kg) 10 days and 2 days before sacrifice. Animals were housed in social groups and maintained in a 12 h:12 h light:dark cycle and had free access to water and diet. At necropsy, blood collection by intracardiac aspiration (∼250 µl) was performed in EDTA-treated tubes. Blood samples were spun at 13,000 *g* for 15 minutes, aliquoted and stored at -80°C until measurement of plasma level of CTX-I and P1NP. Uterus were collected and weighted to ensure optimum ovariectomy. Right and left tibias and right and left femurs were collected and cleaned of soft tissue. Right femurs were wrapped in saline-soaked gauze and frozen at -20°C until use. Left tibias were snap frozen in RNAlater and store at -80°C until use. Other bones were stored in 70% ethanol.

### 2.7. High resolution X-ray microCT

MicroCT analyses were performed at the right tibia with a Bruker 1272 microtomograph operated at 70 kV, 140 μA, 1000 ms integration time. The isotropic pixel size was fixed at 4 μm, the rotation step at 0.25° and exposure was performed with a 0.5 mm aluminum filter. A trabecular volume of interest was located 0.5 mm below the growth plate at the proximal end and extended 2 mm down. A cortical volume of interest was also located at the midshaft tibia and extended 0.5 mm down. All histomorphometrical parameters were measured according to guidelines and nomenclature proposed by the American Society for Bone and Mineral Research ^(30)^.

### 2.8. Bone histomorphometry

After microCT scans, right tibias were embedded undecalcified in pMMA at 4°C. For each animal, four non serial longitudinal sections (∼50 µm apart) were counterstained with calcein blue for dynamic histomorphometry and four additional sections were stained for TRAcP as previously described ^(31)^. Histomorphometrical parameters of bone formation have been computed using the CalceinHisto software developed by Professor Rob van’t Hof (Institute of Ageing and Chronic Disease – University of Liverpool, UK). Standard bone histomorphometrical nomenclatures, symbol and units were used as described in the guidelines of the American Society for Bone and Mineral Research ^(32)^. The identity of the sections was not revealed to the pathologist until the end of all measurements.

### 2.9. Assessment of bone strength

Whole-bone strength of right femurs was assessed by 3-point bending as described previously ^(16,33)^ and in accordance with published guidelines ^(34)^. Three-point bending strength was measured with a constant span length of 10 mm. Bones were tested in the antero-posterior axis with the posterior surface facing upward, centered on the support and the pressing force was applied vertically to the midshaft of the bone. Each bone was tested with a loading speed of 2 mm.min^-1^ until failure with a 500 N load cell on an Instron 5942 device (Instron, Elancourt, France) and the load-displacement curve was recorded at a 100 Hz rate by the Bluehill 3 software (Instron). Ultimate load, ultimate displacement, stiffness and work to fracture were calculated as indicated in ^(35)^. The yield load was calculated with the 0.2% offset method. Post-yield displacement was also computerized.

### 2.10. Bone ECM material evaluation

For collagen crosslink analysis in osteoblast cultures, extracellular matrix was prepared as already reported ^(15)^. Spectral analysis was performed using a Bruker Hyperion 3000 infrared microscope coupled to a Bruker Vertex 70 spectrometer and the standard single element MCT detector. Mid-infrared spectra were recorded at a resolution of 4 cm^-1^ (Spectral range 750–2000 cm^-1^), with 128 accumulations in transmission mode. For analysis of bone specimen, left femurs were embedded undecalcified in pMMA after dehydration and infiltration as previously reported ^(16)^. One micrometer-thick cross-section of the midshaft femur was cut with an ultramicrotome (Leica EM UC7, Leica microsystems, Nanterre, France) and deposited on BaF2 windows. Spectral analysis was performed with the same setup (Bruker Hyperion 3000 infrared microscope/Vertex 70 spectrometer) using a 64 × 64 focal plane array detector. A field of view of 540 × 540 µm covering the posterior quadrant of the femur midshaft was analyzed. Mid-infrared spectra were recorded at a resolution of 8 cm^-1^ (spectral region 900-2000 cm^-1^), with 32 accumulations in transmission mode.

For extracellular matrix and femur cross-section, background spectra were collected under identical conditions from the same BaF_2_ windows at the beginning and end of each experiment to ensure instrument stability. Post-processing was performed using Matlab R2021b (The Mathworks, Natick, CA) and included Mie scattering correction, second derivative spectroscopy and curve fitting routines as previously reported ^(36)^. ECM material parameters analyzed for osteoblast cultures was collagen maturity (area ratio 1660/1690 cm^-1^). Combination indices were computed according to Chou and Talalay method ^(37)^ assuming a bliss independence model as:

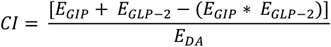

where E_GIP_, E_GLP-2_ and E_DA_ represents the effects on collagen maturity observed with GIP alone, GLP-2 alone or unimolecular dual GIP/GLP-2 analogue, respectively, as compared with vehicle-treated cultures.

Bone ECM material parameters for cortical cross section were: mineralization degree (area of v1,v3 phosphate/area amide I); mineral crystallinity/maturity (intensity ratio 1030 cm^-1^/1020 cm^-1^), carbonate/phosphate ratio (intensity v3 carbonate located at ∼1415 cm^-1^/1030 cm^-1^), trivalent collagen crosslink content (intensity 1660 cm^-1^/area Amide I), divalent collagen crosslink content (intensity 1690 cm^-1^/area Amide I) and collagen maturity (intensity ratio 1660 cm^-1^/1690 cm^-1^) ^(38)^. Histogram distribution for each compositional parameter were fitted with a Gaussian model and considered normally distributed if the R² coefficient was >0.95. In the present study, no histogram deviated from normal distribution. For each of the compositional parameters, the mean of the pixel distribution (excluding non-bone pixels) was computed.

### 2.11. GL-0001 degradation

Fifty micrograms of PYY_1-36_ or GL-0001_1-33_ were incubated at 37°C on a plate shaker in 50 mM triethanolamine/HCl (pH 7.8) with 0.05 U of pure DPP-4 enzyme for 0, 0.5, 1, 2 and 8 hours. Reactions were terminated, as appropriate, via the addition of 10% (v/v) trifluoroacetic acid/water. Reaction mixes were separated by reverse phase high-performance liquid chromatography (RP-HPLC) with absorbance monitored at 214 nm using a Thermo Separation SpectraSYSTEM UV2000 detector. High-performance liquid chromatography peaks were collected and identified via MALDI-ToF MS on a PerSeptive Biosystems Voyager-DE Biospectrometry (Hertfordshire, UK). Percentage of intact and cleaved peptides were computed based on the area of the corresponding peak on HPLC profile.

### 2.12. Statistical analysis

Statistical analyses were performed with GraphPad Prism 8.0 (GraphPad Software, La Jolla, CA, USA). The Brown-Forsythe test of equal variance was applied and when normality was respected, ordinary one-way ANOVA with multiple post hoc Dunnett comparisons were completed. When normality was not respected, Kruskal-Wallis test with multiple post hoc Dunn’s comparisons were computed. Linear regression was computed to correlate biomechanical properties with bone turnover, bone structure and bone ECM material properties. Unless otherwise stated, data are presented as mean ± SD. Differences at p<0.05 were considered significant.

## 3. RESULTS

### 3.1. Co-administration of GIP and GLP-2 agonists modulated bone ECM material properties

As a proof of concept, and to follow our previous work on first generation analogues ^(14,15)^, we co-administered the GIP and GLP-2 analogues, [D-Ala^2^]GIP_1-30_ and [Gly^2^]GLP-2, in murine MC3T3-E1 osteoblast cultures at a concentration of 100 pM. This concentration was previously found to exhibit a pronounced biological response ^(14)^. As compared with vehicle or with single agonist, co-administration resulted in higher levels of intracellular cAMP (Figure 1A). The same pattern was also observed for the expression of lysyl oxidase, a key enzyme involved in enzymatic collagen crosslinking (Figure 1B), and ultimately led to higher levels of collagen crosslinking (Figure 1C). These effects were abolished if osteoblasts were previously treated with 2’,5’-dideoxyadenosine, a selective adenylyl cyclase inhibitor (Figures 1D-E), suggesting that enhanced expression of lysyl oxidase and collagen crosslinking in response to single agonists or co-administration, are cAMP-dependent. The use of *β* -aminopropionitrile (bAPN), a selective lysyl oxidase inhibitor that acts by binding irreversibly to the lysyl oxidase binding site ^(26)^, hampered the response observed with GIP or GLP-2 single agonists and with co-administration (Figures 1F). Together these data supported our initial hypothesis that co-administration of GIP and GLP-2 analogues exert additive effects and enhanced collagen post-processing to a higher extent as seen with single parent peptide, and hence may improve bone ECM material properties through a cAMP-LOX pathway.

**Figure 1:**
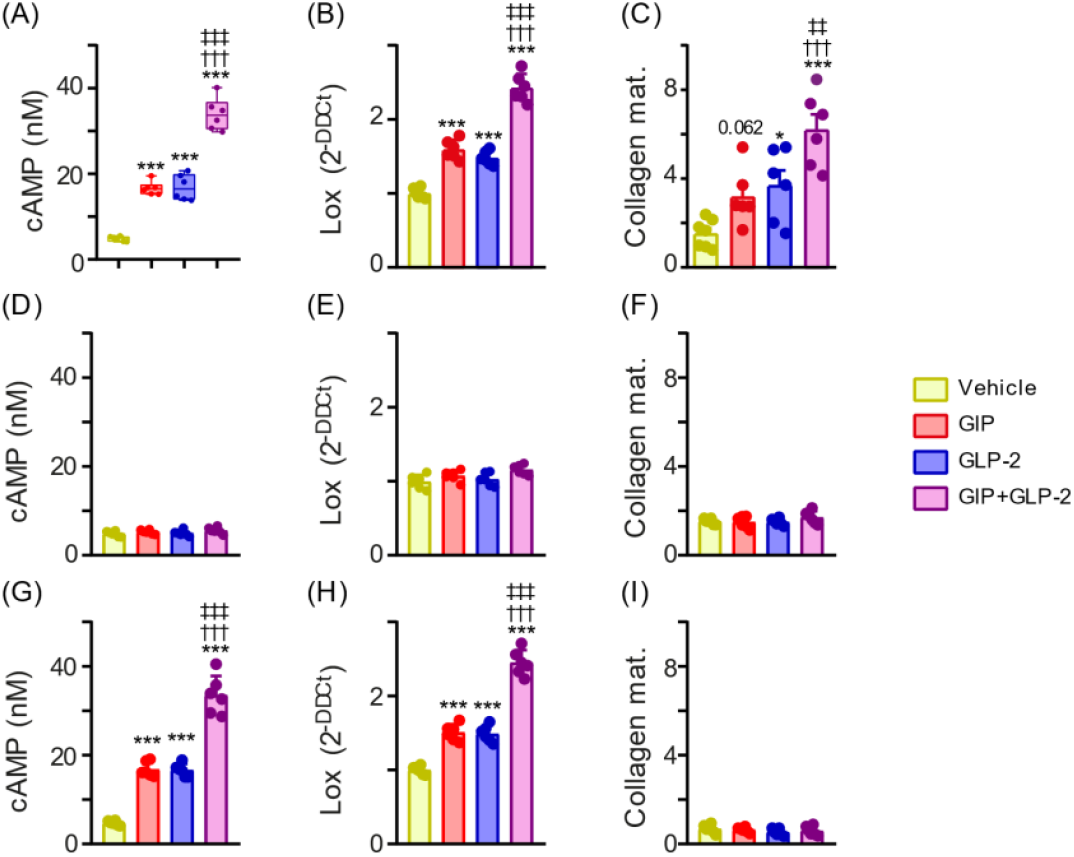
Coadministration of GIP and GLP-2 single analogues enhanced collagen maturity through a cAMP-LOX pathway. Murine MC3T3-E1 osteoblasts were treated with either vehicle, 100 pM [D-Ala^2^]GIP_1-30_, 100 pM [Gly^2^]GLP-2 or co-administration of both analogues. (A, D, G) cAMP levels were determined 48 hrs after treatment by EIA. (B, E, H) Lysyl oxidase expression and (C, F, I) collagen maturity were determined after 14 days of treatment with the above peptides. In order to elucidate the molecular pathway involved in enhanced collagen maturity, MC3T3-E1 cells were either treated with 2’,5’ dideoxyadenosine (D-F) or β-aminopropionitrile (G-I). ^*^: p<0.05 and ^* * *^: p<0.001 vs. vehicle; †††: p<0.001 vs. [D-Ala^2^]GIP_1-30_; ‡‡‡: p<0.001 vs. [Gly^2^]GLP-2.

### 3.2. Development and validation of a series of unimolecular dual GIP/GLP-2 analogues

We then performed a sequence alignment between human GIP and human GLP-2. As the first 30 amino acids of human GIP and GLP-2 are sufficient to induce biological activity ^(39,40)^, we only used this portion for sequence alignment. Human GIP and GLP-2 share ∼67% similarity with divergence at position 7, 12, 13, 18, 19, 28 and 29. The observed similarity and divergence at key position was used as the basis for developing a new class of synthetic molecules, namely unimolecular GIP/GLP-2 dual analogues, discrepancy in pharmacokinetic profiles of GIP and GLP-2 single agonists. Optimization of the consensus sequence was performed in silico based on available information regarding structure-activity relationship and published docking studies between human GIP or GLP-2 and their respective receptors ^(40-46)^. To note, a glycine residue was introduced at position 2 to confer resistance to DPP-4 degradation ^(47)^. We synthesized nine different unimolecular dual GIP/GLP-2 analogues and further investigated their biological potency in vitro.

As binding, but more importantly, activation of cAMP appeared crucial in co-administration studies, we thought to determine whether unimolecular dual GIP/GLP-2 analogues could bind to both the human GIPr and human GLP-2r and activate the production of cAMP (Table 1). Interestingly, in our competition assay, most dual GIP/GLP-2 analogues bound to the human GIPr with affinity, as represented by IC_50_, similar to GIP_1-30_ at the exception of GL-0005, GL-0006 and GL-0009. However, only GL-0001, GL-0007 and GL-0008 significantly raised intracellular levels of cAMP, represented by higher FRET ratio, suggesting binding to and activation of the human GIPr. At the human GLP-2r, GL-0001, GL-0004, GL-0007 and GL-0008 presented with similar binding affinity as compared with human GLP-2. Interestingly, GL-0001, GL-0004 and GL-0007 led to higher intracellular levels of cAMP, suggesting here again that these molecules not only bind to but also activate the human GLP-2r. We also computed changes in FRET ratio as compared with the conventional ligand at both receptors. These data were used to compute a co-agonism score to appreciate activity at both receptors (Table 1). Theoretically, a score superior to 0.5 suggest co-agonism. These data suggest that GL-0008 and GL-0004 are single agonists of the human GIPr and human GLP-2r, respectively. Interestingly, these data also support a co-agonism action of GL-0001 and GL-0007.

**Table 1:**
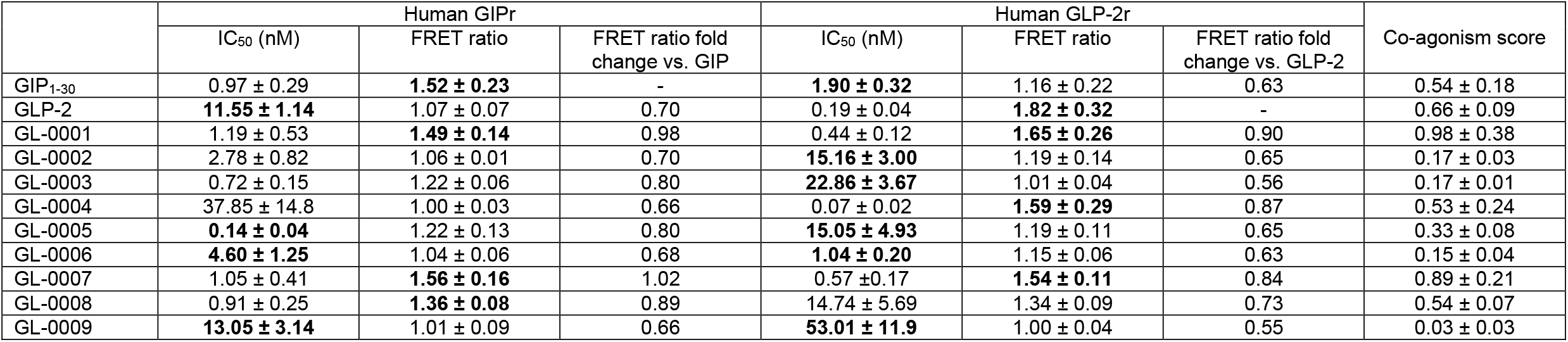
Affinity and potency of unimolecular dual GIP/GLP-2 analogues. Binding to the receptor was investigated in competition with FAM-GIP_1-30_ or FAM-GLP-2 at the human GIPr and the human GLP-2r, respectively. Intracellular levels of cAMP were determined by ratiometric Förster resonance energy transfer (FRET) using the H74 Epac-based FRET biosensor. FRET ratio was investigated at the IC_50_ concentration for all molecules (GIP_1-30_, GLP-2 and dual analogues). FRET ratio was standardized as 1 in the presence of vehicle only. Augmentation in FRET ratio indicates increasing intracellular concentration of cAMP. FRET ratio fold change was computed as FRET ratio of dual analogue divided by FRET ratio of endogenous ligand. Coagonism was calculated as 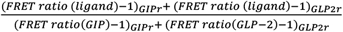. All experiments were carried in duplicate. IC_50_ and FRET ratio are represented as mean ± SEM of three independent experiments. Bold value represents p<0.05 vs. endogenous ligand.

|  | Human GIPr |  |  | Human GLP-2r |  |  | Co-agonism score |
| --- | --- | --- | --- | --- | --- | --- | --- |
|  | IC<sub>50</sub> (nM) | FRET ratio | FRET ratio fold change vs. GIP | IC<sub>50</sub> (nM) | FRET ratio | FRET ratio fold change vs. GLP-2 |  |
| GIP<sub>1-30</sub> | 0.97 ± 0.29 | **1.52 ± 0.23** | - | **1.90 ± 0.32** | 1.16 ± 0.22 | 0.63 | 0.54 ± 0.18 |
| GLP-2 | **11.55 ± 1.14** | 1.07 ± 0.07 | 0.70 | 0.19 ± 0.04 | **1.82 ± 0.32** | - | 0.66 ± 0.09 |
| GL-0001 | 1.19 ± 0.53 | **1.49 ± 0.14** | 0.98 | 0.44 ± 0.12 | **1.65 ± 0.26** | 0.90 | 0.98 ± 0.38 |
| GL-0002 | 2.78 ± 0.82 | 1.06 ± 0.01 | 0.70 | **15.16 ± 3.00** | 1.19 ± 0.14 | 0.65 | 0.17 ± 0.03 |
| GL-0003 | 0.72 ± 0.15 | 1.22 ± 0.06 | 0.80 | **22.86 ± 3.67** | 1.01 ± 0.04 | 0.56 | 0.17 ± 0.01 |
| GL-0004 | 37.85 ± 14.8 | 1.00 ± 0.03 | 0.66 | 0.07 ± 0.02 | **1.59 ± 0.29** | 0.87 | 0.53 ± 0.24 |
| GL-0005 | **0.14 ± 0.04** | 1.22 ± 0.13 | 0.80 | **15.05 ± 4.93** | 1.19 ± 0.11 | 0.65 | 0.33 ± 0.08 |
| GL-0006 | **4.60 ± 1.25** | 1.04 ± 0.06 | 0.68 | **1.04 ± 0.20** | 1.15 ± 0.06 | 0.63 | 0.15 ± 0.04 |
| GL-0007 | 1.05 ± 0.41 | **1.56 ± 0.16** | 1.02 | 0.57 ± 0.17 | **1.54 ± 0.11** | 0.84 | 0.89 ± 0.21 |
| GL-0008 | 0.91 ± 0.25 | **1.36 ± 0.08** | 0.89 | 14.74 ± 5.69 | 1.34 ± 0.09 | 0.73 | 0.54 ± 0.07 |
| GL-0009 | **13.05 ± 3.14** | 1.01 ± 0.09 | 0.66 | **53.01 ± 11.9** | 1.00 ± 0.04 | 0.55 | 0.03 ± 0.03 |

### 3.3. Evaluation of the biological activity of unimolecular dual GIP/GLP-2 analogues

As higher expression of lysyl oxidase and higher collagen maturity were observed in co-administration study, we investigated whether unimolecular dual GIP/GLP-2 analogues could recapitulate these findings. Similar to co-administration study, a dose of 100 pM was used to estimate the potency of unimolecular dual GIP/GLP-2 analogues. Interestingly, at the exception of GL-0002, GL-0003 and GL-0009, all other unimolecular dual GIP/GLP-2 analogues significantly increased *Lox* expression as compared with vehicle-treated murine cultures (Figure 2A). However, as compared with GIP- or GLP-2-treated cultures, only GL-0001 and GL-0007 significantly improved the extent of *Lox* expression (23% - 34%, p<0.001). As a result, unimolecular dual GIP/GLP-2 analogues that showed positive effects on *Lox* expression resulted in higher collagen maturity in murine cultures (Figure 2B). Of note, GL-0001 and GL-0007 were the only two dual molecules that significantly enhanced the extent of collagen crosslinking beyond the effects of GIP or GLP-2.

**Figure 2:**
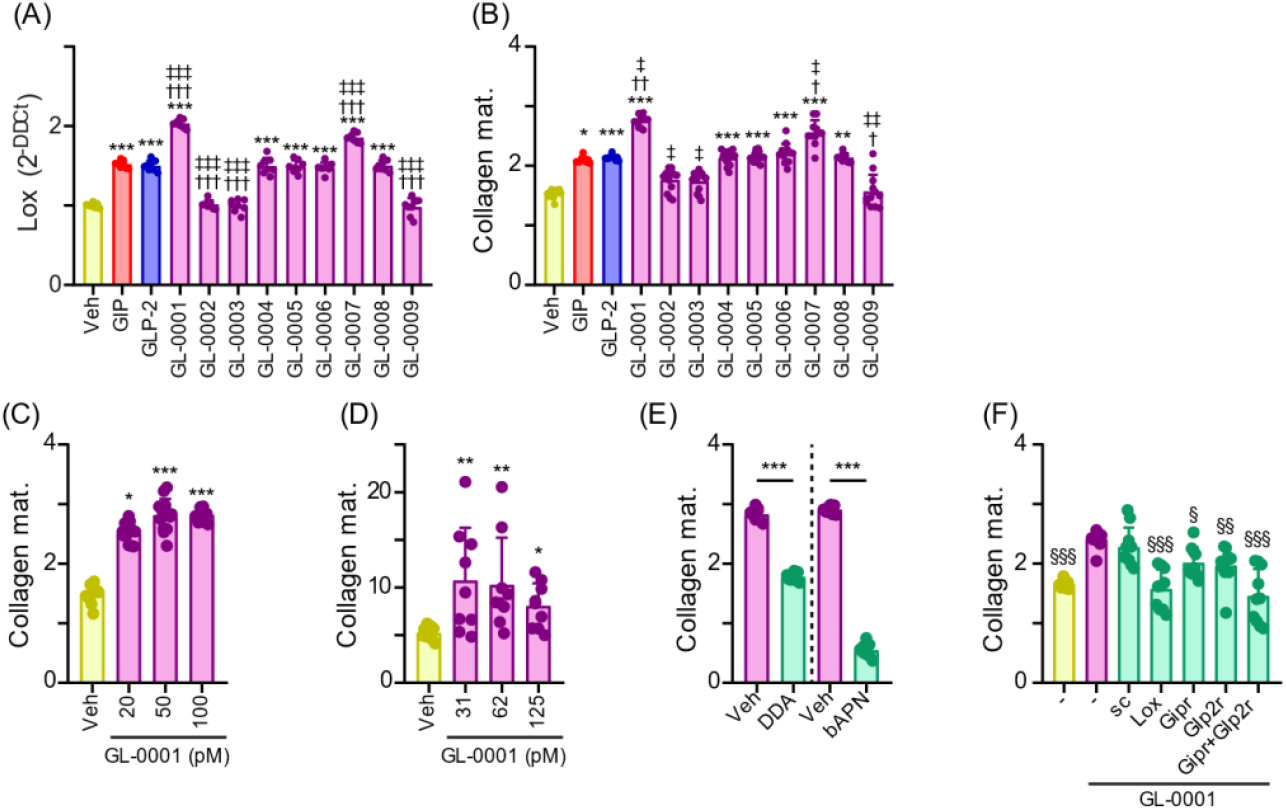
Effects of newly designed unimolecular dual GIP/GLP-2 analogues on lysyl oxidase expression and collagen maturity. (A-B) Murine MC3T3-E1 osteoblasts were treated with either vehicle, 100 pM [D-Ala^2^]GIP_1-30_, 100 pM [Gly^2^]GLP-2 or 100 pM of unimolecular dual GIP/GLP-2 analogues. Lysyl oxidase expression and collagen maturity were determined after 14 days of treatment with the above peptides. (C) Murine MC3T3-E1 osteoblasts were treated with either vehicle or several concentrations (20 – 100 pM) of GL-0001 prior collagen maturity evaluation. (D) Human primary osteoblasts cells were cultured in the presence of several concentrations (31 – 125 pM) of GL-0001 prior collagen maturity evaluation. (E) MC3T3-E1 cells were either treated with 2’,5’ dideoxyadenosine (DDA) or β-aminopropionitrile (bAPN) in the presence of 100 pM GL-0001 prior collagen maturity evaluation. (F) MC3T3-E1 cells were incubated with 100 pM GL-0001 and siRNA targeting lysyl oxidase (Lox), GIPr (Gipr), GLP-2r (Glp2r), both receptors (Gipr+Glp2r) or a scrambled siRNA (sc). Control cells were treated in the absence of siRNA (-). ^*^: p<0.05, ^* *^:p<0.01 and ^* * *^: p<0.001 vs. vehicle; †: p<0.05, ††: p<0.01 and †††: p<0.001 vs. [D-Ala^2^]GIP_1-30_; ‡: p<0.05, ‡‡: p<0.01 and ‡‡‡: p<0.001 vs. [Gly^2^]GLP-2. §: p<0.05, §§: p<0.01 and §§§: p<0.001 vs. cells treated with GL-0001 in the absence of siRNA.

We next thought to decipher whether the action of unimolecular dual GIP/GLP-2 analogues were due to synergism at both the GIPr and GLP-2r. In our case, it was not possible to perform the conventional isobologram analysis approach as GIP/GLP-2 co-agonism is due to a single molecule, not a combination of two or more molecules. However, we computed the combination index according to Chou & Talalay ^(37)^, assuming a bliss independence model. Effects on collagen maturity in murine cultures was used for combination index determination. Interestingly, GL-0001 and GL-0007 exhibited a combination index of 0.46 ± 0.07 and 0.65 ± 0.07, respectively, suggesting that both molecules possess synergism properties (Supplementary Table 2). GL-0005, GL-0006 and GL-0008 presented with combination index suggesting additive effects. On the other hand, GL-0002, GL-0003, GL-0004 and GL-0009 presented with combination index indicative of moderate to very strong antagonism.

In addition, improvement of collagen maturity in murine cultures observed with GL-0001 was not restricted to a dose of 100 pM but was already achieved at a lower dose of 20 pM (Figure 2C). Recent reports have suggested that activities at the murine and human receptors were not identical at the level of pancreatic islet and this finding needed to be extended to bone receptors ^(48)^. As such, we thought to assess whether positive action of GL-0001 was restricted to rodent or whether it could be transferable in a human context. We treated primary human osteoblast cultures with ascending dose of GL-0001 ranged 31-125 pM (Figure 2D). GL-0001 significantly enhanced the extent of collagen crosslinking with maximum effects already encountered at 31 pM, suggesting that GL-0001 was also capable of activating rodent and human receptors.

We next tried to decipher whether the observed effects on collagen maturity was due to activation of the cAMP-Lox pathways as seen for co-administration. Murine osteoblast cultures were repeated in the presence of the selective adenylyl cyclase inhibitor 2’,5’-DDA. Here again, the effects of GL-0001 were blocked by pretreatment with this pharmacological inhibitor suggesting that GL-0001 requires a functional adenylyl cyclase to exert their effects (Figure 2E). Similarly, pre-treatment with bAPN significantly blocked the action of GL-0001 on collagen maturity supporting the involvement of lysyl oxidase in this process. Furthermore, use of specific siRNA targeting Lox, Gipr, Glp2r or both receptors resulted in lower collagen maturity as compared with absence of siRNA suggesting that these three genes are required for GL-0001 effects on collagen maturity (Figure 2F).

### 3.4. Introduction of Gly² demonstrate resistance to dipeptidyl peptidase-4 degradation

We next verified whether the presence of a glycine residue at position 2 confers resistance to DPP-4 as expected. PYY_1-36_ was used as a positive control. As reported in Supplementary Figure 1, for PYY_1-36_, DPP-4 degradation was represented by occurrence of the PYY_3-36_ fragment peptide, consistent with previous literature ^(49)^. For GL-0001, the 3-33 fragment occurred only at 8h, suggesting Gly2 confers resistance to DPP-4. Indeed, the resistance half-life of GL-0001 was estimated at ∼20 h as compared with ∼8.5 minutes for PYY_1-36_. However, as GL-0001_1-33_ could be degraded, we next ascertain whether GL-0001_3-33_ was potent in modulating collagen maturity in vitro. Interestingly, the cleaved form of GL-0001 did not induce a change in the collagen maturity response as compared with vehicle and as opposed to the intact form. More importantly, when both forms, intact and cleaved, were co-added in the cultures, it did not reduce the extent of collagen maturity as compared with the intact form only.

### 3.5. First-in-class unimolecular dual GIP/GLP-2 analogue GL-0001 improved bone strength and bone microstructure in ovariectomy-induced bone fragility

We decided to further evaluate the benefit of unimolecular dual GIP/GLP-2 analogue in the OVX-induced bone fragility model with zoledronic acid as a positive comparator. Efficiency of ovariectomy has been assessed post-mortem by uterus mass (Supplementary Figure2). None of the pharmacological interventions influenced either body weight or blood glucose levels (Supplementary Figure 2). We investigated the biomechanical resistance of long bones following saline or treatment administration. Interestingly, in the load-deformation curves, one can notice that deformation is reduced in vehicle-treated OVX mice suggesting that macroscopical fracture occurs rapidly after coalescence of microcracks (Figure 3A). This feature is not ameliorated by zoledronic acid administration. However, administration of GL-0001 or GL-0007 restored this pattern. Interestingly, GL-0001, but neither GL-0007 nor zoledronic acid, significantly improved ultimate load by 9% (p=0.021), post-yield deformation by 70% (p=0.013) and energy-to-fracture by 37% (p=0.008) (Figure 3B). More importantly, the effects on ultimate load and energy-to-fracture were significantly different as compared with zoledronic acid.

**Figure 3:**
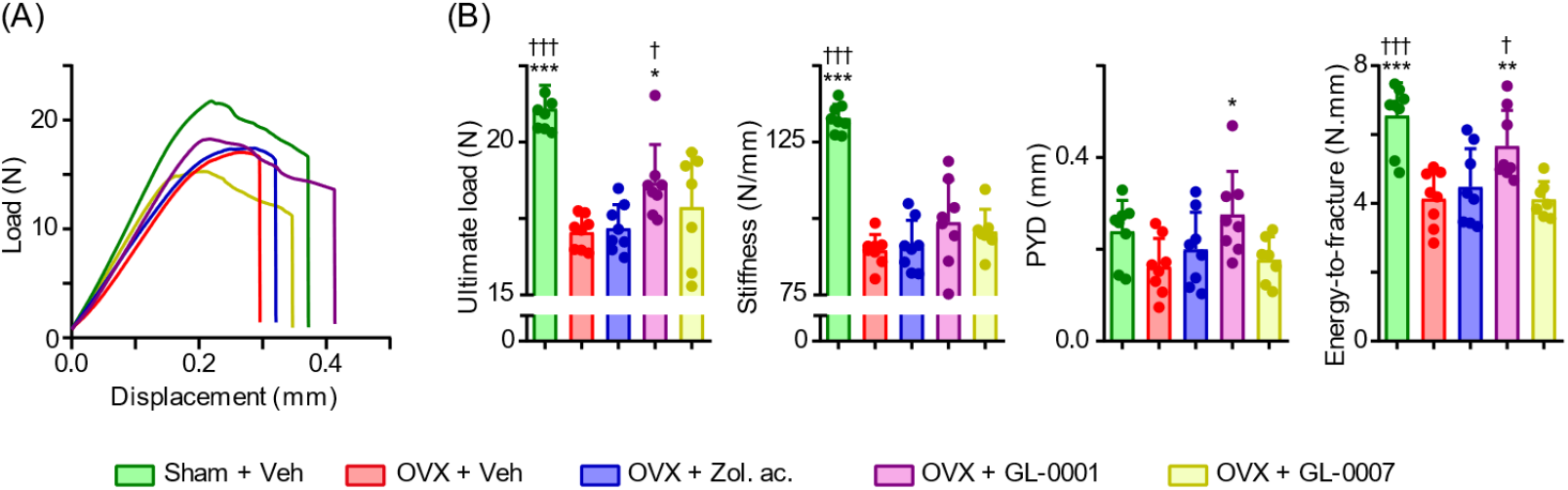
Administration of GL-0001 improved bone biomechanical response in OVX mice. OVX mice were administered with either vehicle, 25 nmoles/kg/day GL-0001, 25 nmoles/kg/day GL-0007 or once iv 100 µg/kg zoledronic acid. (A) Representative load/displacement curves obtained after 8 weeks of treatment and (B) Biomechanical parameters including ultimate load, stiffness, post-yield displacement (PYD) and energy-to-fracture. ^*^: p<0.05, ^* *^: p<0.01 and ^* * *^: p<0.001 vs. OVX+vehicle; †: p<0.05 and †††: p<0.001 vs. OVX+zoledronic acid.

We next investigated whether improvement in the biomechanical response of bone following GL-0001 administration was due to changes in microstructure. As shown in Figures 4A-B, vehicle-treated OVX animals presented with a non-significant reduction in trabecular bone mass, due to a lower number of trabeculae, but also owing to a slight and non-significant reduction in cortical thickness at the mid tibial diaphysis. As expected, zoledronic acid administration significantly enhanced trabecular bone mass by 85% (p<0.001), trabeculae numbers by 106% (p<0.001) and cortical thickness by 6% (p=0.037). Interestingly, GL-0001, but not GL-0007, significantly increased trabecular bone mass by 49% (p=0.009) and trabecular thickness by 15% (p=0.001) but had no significant effects on trabeculae numbers and cortical thickness. To better apprehend the cellular mechanism linked to observed changes in microstructure, we evaluated the circulating levels of PTH_1-84_, a key hormone in calcium and phosphate metabolism, and CTX-I and P1NP, two key molecules relevant to bone remodeling. No changes in circulating levels of PTH_1-84_ were observed between the experimental groups (Figure 4C). As expected, circulating levels of the bone resorption marker CTX-I was significantly elevated by 41% (p<0.001) in vehicle-treated OVX animals (Figure 4C). Interestingly, GL-0001-, GL-0007- and zoledronic acid-treated OVX mice presented with significant reductions in this parameter by 31% (p<0.001), 19% (p<0.001) and 24% (p<0.001), respectively (Figure 4C). At the cellular level, the number of osteoclasts per bone perimeter and bone surfaces covered with osteoclast showed similar patterns as compared with CTX-I (Figures 4D). Zoledronic acid also led to a significant reduction in the circulating levels of the bone formation marker P1NP (−25%, p=0.011) whilst no changes were observed with either GL-0001 or GL-0007 (Figure 4E). Dynamic bone histomorphometrical parameters suggested a reduction in active mineralizing surface and in bone formation rate with zoledronic acid but not with either GL-0001 or GL-0007 (Figure 4E).

**Figure 4:**
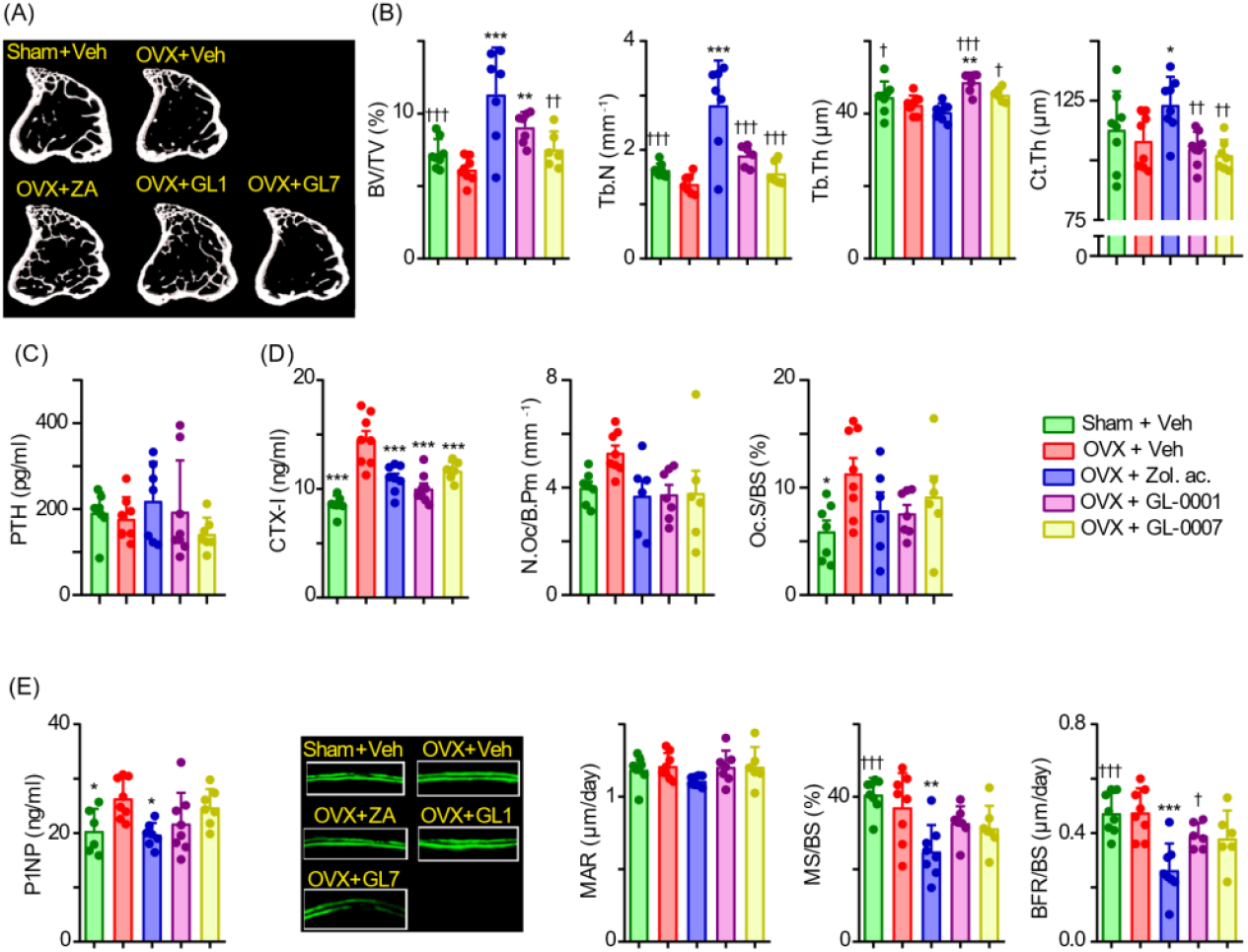
Histomorphometrical analyses of the appendicular skeleton in OVX mice. (A) Representative 3D models of the tibia proximal metaphysis and (B) trabecular and cortical microarchitecture parameters. (C) Circulating levels of parathyroid hormone (PTH). (D) Circulating levels of the resorption marker CTX-I and static osteoclast histomorphometry parameters. (E) Circulating levels of the bone formation marker P1NP and dynamic histomorphometry parameters. ^*^: p<0.05, ^* *^: p<0.01 and ^*^ * *: p<0.001 vs. OVX+vehicle; †: p<0.05, ††: p<0.01 and †††: p<0.001 vs. OVX+zoledronic acid.

Although, GL-0001 has not been designed to reduce osteoclast formation and/or resorption, the observed reduction in CTX-I levels and the apparent trend in lowering the number of osteoclasts and osteoclast surfaces, led us to investigate whether co-administration of GIP and GLP-2 or GL-0001 had anti-osteoclastic properties. When single GIP and GLP-2 agonists were co-administered to murine osteoclast precursors cells, they significantly reduced the extent of osteoclast formation as compared with vehicle or single agonists (Supplementary Figure 3A). GL-0001 administration in murine osteoclast precursors resulted in the same pattern (Supplementary Figure 3B). This effect was also seen in human osteoclast precursor cultures, where GL-0001 dose-dependently reduced osteoclast formation (Supplementary Figure 3C). More importantly, in human osteoclast cultures, GL-0001 dose-dependently reduced osteoclast resorption (Supplementary Figure 3D).

### 3.6. First-in-class unimolecular dual GIP/GLP-2 analogue GL-0001 improved bone ECM material properties in ovariectomy-induced bone fragility

As unimolecular dual GIP/GLP-2 analogues have been developed to improve bone ECM material properties and do so in vitro, we thought to determine whether such effects were preserved in the OVX-induced bone fragility mouse model. As presented in Figure 5A, administration of either GL-0001 or GL-0007 did not modify either *Col1a1* or *Plod2* expression. However, GL-0001, but not GL-0007, significantly downregulated by 28% the expression of alkaline phosphatase (p<0.001) and upregulated by 65% the expression of lysyl oxidase (p=0.028). We next determined whether changes in *Alpl* and *Lox* expression, two key molecules involved in bone ECM maturation and mineralization, were related to modifications of bone ECM material properties. Fourier transform infrared imaging highlighted changes in mineralization degree and collagen maturity over the full cortical width (Figure 5B). Indeed, when quantified, the degree of mineralization of the bone ECM was significantly reduced by 2% (p<0.001) in GL-0001-treated OVX mice as compared with vehicle-treated OVX mice. Similarly, collagen maturity was enhanced in GL-0001-treated OVX mice by 10% (p=0.011). None of these effects were observed with either zoledronic acid or GL-0007 (Figure 5C). To further understand the contribution of mature and immature collagen crosslinks to the observed change in collagen maturity, we monitored their spectral signature in the same bone sample (Figure 5C). We evidenced that mature crosslinks and immature crosslinks were increased by 3% (p=0.043) and reduced by 8% (p<0.001), respectively suggesting an acceleration in the maturation of collagen post-processing (Figure 5C). None of the other ECM material properties were altered in any of the studied groups.

**Figure 5:**
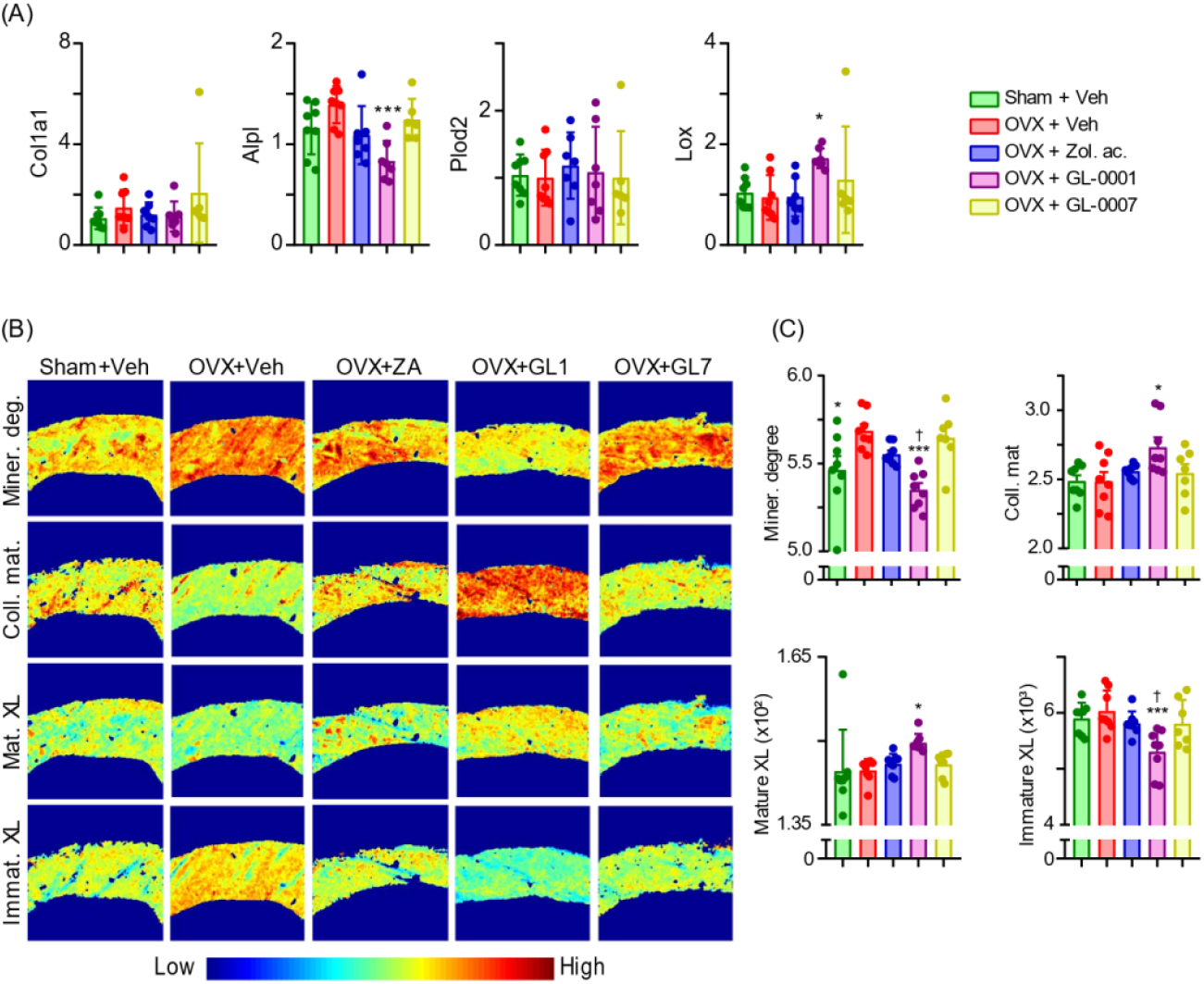
GL-0001 ameliorated bone ECM material properties. (A) Expression levels of gene involved in post-translational modifications of bone ECM. (B) Representative Fourier transform infrared imaging (FTIRI) images and (C) quantitative FTIRI parameters. Mineralization degree (Miner. Degree) represents the area ratio of the v1,v3 PO4/Amide, collagen maturity was computed as the ratio between trivalent mature crosslink (mature XL)/divalent immature crosslink (immature XL). Mature and immature crosslinks were normalized by Amide I area. ^*^: p<0.05 and ^* * *^: p<0.001 vs. OVX+vehicle; †: p<0.05 vs. OVX+zoledronic acid.

Finally, in order to better understand whether changes at the microstructure and/or material levels were linked to the positive effects observed at the biomechanical level, we performed a linear regression analysis between CTX-I level, cortical thickness, collagen maturity, mineralization degree and the energy-to-fracture. Interestingly, CTX-I level (R² = 0.16, p=0.13) and cortical thickness (R² = 0.01, p=0.71) were not significantly correlated to energy-to-fracture. However, amelioration of collagen maturity (R² = 0.72, p<0.001) and possibly lower mineralization degree (R² = 0.24, p=0.052) seemed linked to improvement in bone biomechanical resistance, suggesting that targeting bone ECM material properties could enhance bone resistance to fracture (Figure 6).

**Figure 6:**
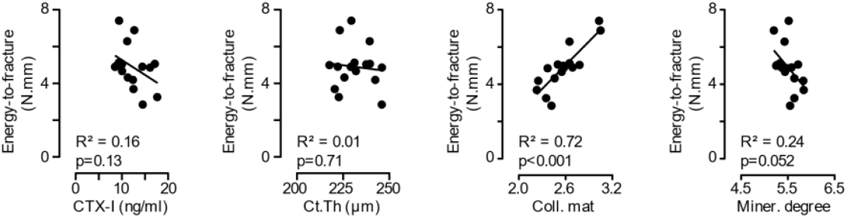
Regression analyses between biomechanical, CTX-I levels, cortical thickness, and bone ECM material properties. Linear regressions were performed with GraphPad Prism and R^2^ and p values are reported.

## 4. DISCUSSION

All current therapeutic approaches to reduce the burden of fragility fracture rely on increasing bone mineral density as a surrogate to enhance bone strength. In contrast, based on our previous work with single parent peptides ^(13-21)^, we conceived and reported herein the development of a first-in-class unimolecular dual GIP/GLP-2 analogues, GL-0001, establishing it as more efficacious than zoledronic acid in reducing bone fragility in the OVX-mice model of bone fragility. Through a series of chemical changes that yielded a hybrid peptide sequence, we identified GL-0001 as a high-potency balanced dual analogue capable to bind to and activate the GIPr and GLP-2r, resistant to DPP-4 degradation, that enhances collagen maturity in vitro in murine and human osteoblast cultures, and improves bone strength mostly by modulating bone ECM material properties in vivo.

Interestingly, GL-0001 also limited bone osteoclast resorption and ultimately led to preservation of trabecular bone mass at the appendicular skeleton. Of note, we evidenced that GL-0001 was capable of reducing osteoclast formation but also osteoclast-mediated resorption in osteoclast cultures. Previously, a similar effect was encountered with single GIP analogues ^(50-52)^ or single GLP-2 analogues ^(17,53)^. Furthermore, subcutaneous administration of GIP or GLP-2, or co-administration of GIP and GLP-2 in healthy individuals resulted in a marked decreased in CTX-I levels and were mirrored with an acute suppression of PTH secretion ^(25,54)^. Both receptors appear to be expressed with the same magnitude in parathyroid gland tissues, but only GLP-2, and not GIP, exert antiresorptive actions through reduced secretion of PTH ^(24)^. Currently, function of the GIPr on parathyroid gland remains to be fully elucidated. Interestingly, in our study, we failed to demonstrate any significant change in circulating PTH levels following GL-0001 administration, despite clear reduction in CTX-I levels, ruling out any PTH-mediated effects to explain the anti-resorptive properties of GL-0001. However, we clearly showed that GL-0001 reduced the number of newly generated osteoclasts in murine and in human cultures and also reduced the extent of bone resorption suggesting that the observed reduction in CTX-I levels was more likely due to direct effects on osteoclast precursors/mature osteoclasts. Nevertheless, recent evidence also suggests that joint administration of GIP and GLP-2 in post-menopausal subjects is more potent in reducing circulating levels of CTX-I as compared with single infusion of GIP or GLP-2 ^(55)^. Our data are in accordance with such mechanisms. Indeed, administration of GL-0001 to cultured human osteoclast precursor elicited a significant reduction in osteoclast-mediated bone resorption and osteoclast precursor differentiation. This is also in agreement with the recent observation that GIPr and GLP-2r are expressed by human osteoclasts ^(24)^. However, in our study, despite significant reduction in CTX-I levels with GL-0007 treatment, no effects of this molecule were observed in modulating bone ECM material properties and enhancing bone mechanical resistance in vivo. Our regression study unambiguously evidenced that the improved biomechanical response of bone with GL-0001 was mostly due to changes in bone ECM material properties but not with either cortical thickness or CTX-I levels. Although it is tempting and practical to monitor the effects of unimolecular dual GIP/GLP-2 analogues with a simple blood test in humans, it is not a proof of efficacy in improving bone biomechanical response.

Unimolecular dual GIP/GLP-2 analogues have been developed to bring to the therapeutic arsenal against bone fragility, new molecules that could directly target bone ECM material properties. Single GIP or GLP-2 analogues have been characterized in the past and several analogues exhibit significant effects on bone material properties by targeting lysyl oxidase expression and/or posttranslational modifications of bone collagen including crosslinking and maturity ^(13,17-22,56,57)^. During optimization of the consensus sequence, we noticed that amino acid positions 7, 10, 12 and 13 were crucial to balance the activity of unimolecular GIP/GLP-2 analogues at the GIPr or GLP-2r. An isoleucine at position 12 had been shown previously to be vital for selective GIP activity in a series of unimolecular dual GIP/GLP-1 agonists ^(42)^, whereas substitution of Thr^12^ by Ala^12^ in GLP-2 reduced generation of cAMP ^(41)^. Interestingly, GL-0001 and GL-0007 revealed that dual agonism at the hGIPr and hGLP-2r differs only at positions 10, 31, 32 and 33. The methionine residue at position 10 is the natural amino acid in human GLP-2. In contrast, at this position, human GIP exhibits a tyrosine residue. The use of Met^10^ in GL-0001 conserved activity at both hGIPr and hGLP-2r. The use of Phe^10^ in GL-0007, with similar side chain length as compared with endogenous Tyr^10^, also led to balanced activity at both receptors, at least in vitro. It is not clear now whether bias towards either the GIPr or GLP-2r, as seen for unimolecular dual GIP/GLP-1 compound tirzepatide ^(58)^, could be beneficial in improving bone strength to a greater extent than seen with GL-0001. However, this should be further investigated in the future.

Recently, others have reported the development of another unimolecular dual GIP/GLP-2 analogue, named peptide 11 that shows ∼79% similarity with GL-0001 ^(54)^. However, and in opposition to GL-0001, peptide 11 demonstrated a specie-dependency bias towards either the GIPr or the GLP-2r in human and rodent systems, respectively. This behavior was not found with GL-0001 as similar enhancement of collagen maturity were observed in human and rodent cultures. Furthermore, and to the best of our knowledge, no bone data, have been reported for peptide 11 and as such it is not possible to compare the activity of GL-0001 to this other unimolecular dual GIP/GLP-2 analogue. Nevertheless, GL-0001 lead to significant amelioration of biomechanical resistance to fracture and clearly reversed compromised bone ECM material properties in OVX animals.

In conclusion, we developed a series of unimolecular dual GIP/GLP-2 analogues with the first-in-class molecule, GL-0001, capable of enhancing collagen maturity and directly improving bone biomechanical response and resistance to fracture in vivo. GL-0001, and more broadly unimolecular dual GIP/GLP-2 analogues, represent a new therapeutic solution for the treatment and prevention of bone fragility that targets bone material properties rather than bone mineral density. This innovative pathway represents an interesting and complementary way to the conventional therapeutical arsenal in order to treat bone fragility patients.

## ABBREVIATION LIST

2’,5‘-DDA: 2’, 5’ dideoxyadenosine;
αMEM: alpha minimum essential medium;
ANOVA: Analysis of variance;
bAPN: beta-aminopropionitrile;
BSA: Bovine serum albumin;
CTX-I: C-terminal telopeptide of collagen type I;
cDNA: Complementary desoxyribonucleic acid;
cAMP: Cyclic adenosine monophopshate;
DPP-4: dipeptidyl peptidase-4;
ECM: Extracellular matrix;
EIA: Enzyme immunoassay;
FBS: Fetal bovine serum;
FRET: Förster resonance energy transfert;
GIP: glucose-dependent insulinotropic polypeptide;
GIPr: glucose-dependent insulinotropic polypeptide receptor;
GLP-2: glucagon-like peptide-2;
GLP-2r: glucagon-like peptide-2 receptor;
HEPES: 4-(2-hydroxyethyl)-1-piperazineethanesulfonic acid;
HPLC: High pressure liquid chromatography;
IBMX: 3-isobutyl-1-methylxanthine;
LOX: Lysyl oxidase;
MALDI-ToF MS: Matrix Assisted Laser Desorption Ionization - time-of-flight mass spectrometry;
M-CSF: Macrophage-colony stimulating factor;
MCT: Mercury Cadmium Telluride;
MicroCT: Microcomputed tomography;
OVX: ovariectomy;
P1NP: N-terminal propeptide of type I collagen;
PBMC: Peripheral blood mononuclear cells;
PBS: Phosphate buffer saline;
pMMA: polymethylmethacrylate;
PTH: Parathyroid hormone;
PYY: Peptide Y Y;
qPCR: quantitative polymerase chain reaction;
RIPA: radioimmunoprecipitation assay;
RNA: Ribonucleic acid;
siRNA: Small interfering ribonucleic acid;
sRANKL: soluble receptor activator of nuclear factor κB ligand;
TRAcP: Tartrate resistant acid phosphatase.

## 5. AUTHOR CONTRIBUTION

**Benoit Gobron:** Investigation, Formal analysis, Writing - Original Draft; **Malory Couchot:** Investigation, Formal analysis, Writing – Review & Editing; **Nigel Irwin:** Investigation, Formal analysis, Writing - Review & Editing; **Erick Legrand:** Conceptualization, Resources, Writing - Review & Editing; **Béatrice Bouvard:** Conceptualization, Resources, Writing - Review & Editing, Supervision; **Guillaume Mabilleau:** Investigation, Formal analysis, Writing - Review & Editing, Supervision, Funding acquisition

## 6. ACKNOWLEDGEMENTS

The authors are grateful to Dr Boni (Lentivec platform, University of Angers) for his help in the cloning and preparation of plasmid encoding the human GIPr and GLP-2r, and Ms Mieczkowska (HiMolA Platform, University of Angers) for her help in Fourier transform microspectroscopy. We are thankful to Prof Peter Gardner (University of Manchester) for supplying the Mie scattering correction routine for Matlab and Prof Rob van’t Hof for supplying the CalceinHisto software. Part of this work was supported by a grant from the SATT Ouest valorization (grant number DV2541).

## 8. DATA AVAILABILITY STATEMENT

All relevant data are available from the authors upon reasonable request.

