## Supplementary material for "Development of a first-in-class unimolecular dual GIP/GLP-2 analogue, GL-0001, for the treatment of bone fragility"

#### SUPPLEMENTARY FIGURES

**Supplementary figure 1: Degradation by-products of GL-0001 and biological activity. (**A) HPLC profile of PYY1-36, used as a positive control, and GL-0001 at 0h and 8h post incubation with DPP-4. (B) Intact peptide profile obtained from HPLC profile. (C) Effects of intact and cleaved GL-0001 on collagen maturity and (D) normalized enzymatic collagen crosslink content. *: $p<0.05$ and **: $p<0.01$ vs. vehicle-treated cultures.

**Supplementary figure 2: Body weight, uterus mass and blood glucose in OVX-treated mice.** *: $p<0.05$, **: $p<0.01$ and ***: $p<0.001$ vs. OVX+Veh animals.

**Supplementary figure 3: Effects of co-administration of [D-Ala²]GIP1-30 and [Gly²]GLP-2 or GL-0001 on murine and human osteoclast formation and resorption activity.** (A) Co-administration of GIP and GLP-2 analogues was evaluated in murine Raw264.7 cells treated with soluble RANKL. (B) Effects of GL-0001 on osteoclast formation in murine Raw264.7 cultures. Effects of GL-0001 on (C) osteoclast formation and (D) osteoclast resorption in human PBMCs cultures. *: $p<0.05$, **: $p<0.01$ and ***: $p<0.001$ vs. vehicle; †††: $p<0.001$ vs. [D-Ala²]GIP$_{1\text{-}30}$; ‡‡‡: $p<0.001$ vs. [Gly²]GLP-2.

#### Supplementary Tables

##### Supplementary Table 1: Key resources used in this project

| REAGENT or RESOURCE | SOURCE | IDENTIFIER |
|---|---|---|
| Biological sample | | |
| Fetal bovine serum | Pan Biotech | Cat#P30-1902 |
| Chemicals, peptides, and recombinant proteins | | |
| GIP1-30 | Genecust | N/A |
| GLP-2 | Genecust | N/A |
| GL-0001 | Genecust | N/A |
| GL-0002 | Genecust | N/A |
| GL-0003 | Genecust | N/A |
| GL-0004 | Genecust | N/A |
| GL-0005 | Genecust | N/A |
| GL-0006 | Genecust | N/A |
| GL-0007 | Genecust | N/A |
| GL-0008 | Genecust | N/A |
| GL-0009 | Genecust | N/A |
| Fam-[D-Ala$^2$]GIP$_{1-30}$ | Genecust | N/A |
| Fam-[Gly$^2$]-GLP-2 | Genecust | N/A |
| Soluble human M-CSF | Bio-Techne | Cat#216-MC-025 |
| Soluble human RANKL | Bio-Techne | Cat#390-TN-010 |
| PYY$_{1-36}$ | EZBiolab Ltd | N/A |
| Dipeptidyl peptidase-4 | Sigma-Aldrich | Cat#317640-M |
| Zoledronic acid | Tocris | Cat#6111 |
| 2’,5’dideoxyadenosine | Sigma-Aldrich | Cat#288104 |
| Beta-aminopropionitrile fumarate | Sigma-Aldrich | Cat#A3134 |
| 3-isobutyl-1-methylxanthine (IBMX) | Sigma-Aldrich | Cat#410957 |
| Bovine serum albumin | Sigma-Aldrich | Cat#A4503-100G |
| Nucleozol | Macherey-Nagel | Cat#740404.200 |
| Nucleospin RNA column | Macherey-Nagel | Cat#740406.50 |
| RNAlater | Sigma-Aldrich | Cat#R0901 |
| Critical commercial assays | | |
| Clonetics$^{TM}$ OGM$^{TM}$ osteoblast growth medium | Lonza | Cat#CC-3207 |
| Cyclic AMP EIA kit | Bio-Techne | Cat#KGE002B |
| Maxima first strand cDNA synthesis kit | Thermofisher scientific | Cat#K1672 |
| Taqman Fast advanced master mix | Thermofisher scientific | Cat#4444557 |
| RatLaps CTX-I EIA kit | ImmunoDiagnostic Systems (IDS) | Cat# AC-06F1 |
| Rat/mouse P1NP EIA kit | ImmunoDiagnostic Systems (IDS) | Cat# AC-33F1 |
| Experimental models: Cell lines | | |
| Mouse: MC3T3-E1, subclone 4 | ATCC | Cat#CRL-2593; RRID: CVCL_5440 |
| Mouse: Raw 264.7 cells | ATCC | Cat#TIB-71; RRID: CVCL_0493 |
| Hamster: CHO-K1 cells | ATCC | Cat#CCL-61; RRID: CVCL_0214 |
| Human: Primary osteoblasts | Lonza | Cat#CC-2538 |
| Human: Peripheral blood mononuclear cells | Etablissement Francais du Sang | N/A |
| Experimental models: Organisms/strains | | |
| Mouse: BALB/cJRj | Janvier Labs | |

| | | |
|---|---|---|
| **Oligonucleotides** | | |
| siRNA targeting murine GIPr | Thermofisher scientific | Assay ID s233873 |
| siRNA targeting murine GLP-2r | Thermofisher scientific | Assay ID s211996 |
| siRNA targeting murine lysyl oxidase | Thermofisher scientific | Assay ID s69290 |
| Control scrambled siRNA | Thermofisher scientific | Cat#4390843 |
| Taqman gene expression assay for Lox | Thermofisher scientific | Assay ID: Mm00495386_m1 |
| Taqman gene expression assay for Col1a1 | Thermofisher scientific | Assay ID: Mm00801666_g1 |
| Taqman gene expression assay for Alpl | Thermofisher scientific | Assay ID: Mm00475834_m1 |
| Taqman gene expression assay for Plod2 | Thermofisher scientific | Assay ID: Mm00478767_m1 |
| Taqman gene expression assay for B2m | Thermofisher scientific | Assay ID: Mm00437762_m1 |
| **Recombinant DNA** | | |
| hGip-R | Addgene | Plasmid #14942 |
| cDNA of human GLP-2r | Harvard Plasmid information database (PlasmID) | Plasmid #HsCD00346244 |
| Epac-S-H74 | Addgene | Plasmid #170338 |
| **Software and algorithms** | | |
| NRecon reconstruction software | Bruker | Version 1.6.10.2 |
| Dataviewer software | Bruker | Version 1.5.6.2 |
| CTan software | Bruker | Version 1.20.8.0 |
| Opus 6.5 Software | Bruker | N/A |
| Matlab R2021b | The Mathworks | N/A |
| CalceinHisto software | Supplied by Prof Rob van't Hof | N/A |
| Bluehill 3 Software | Instron | N/A |
| Graphpad Prism 8.0 | GraphPad Software | N/A |
| **Other** | | |
| Lipofectamine RNAimax | Thermofisher scientific | Cat#13778-075 |
| Lipofectamine 3000 | Thermofisher scientific | Cat#L3000-08 |
| Cell strainer | Sarstedt | Cat#833945040 |
| SpectraMax® M2 Microplate Reader | Molecular devices | N/A |
| Alzet osmotic minipupmp | Alzet | Model 1004 |
| Standard rodent diet | Safe | Diet A04 |
| C-18 analytical column | Phenomenex | Cat#00G-3050-E0 |
| Spectra SYSTEM UV2000 | Thermo Separation | N/A |
| Voyager-DE Biospectrometry | PerSeptive Biosystems | N/A |
| Vertex 70 spectrometer | Bruker | N/A |
| Hyperion 3000 infrared microscope | Bruker | N/A |
| 1272 microCT | Bruker | N/A |
| Instron 5942 mechanical device | Instron | N/A |

**Supplementary Table 2: Combination index based on collagen maturity effects.**

| Compound | Combination index | p value | Interpretation |
|---|---|---|---|
| **GL-0001** | 0.46 ± 0.07 | <0.0001 | Synergism |
| **GL-0002** | 1.38 ± 0.25 | 0.0083 | Moderate antagonism |
| **GL-0003** | 4.27 ± 2.06 | <0.0001 | Strong antagonism |
| **GL-0004** | 1.39 ± 0.30 | 0.0004 | Moderate antagonism |
| **GL-0005** | 11.19 ± 33.59 | 0.2846 | Additive |
| **GL-0006** | 0.95 ± 0.12 | 0.3748 | Additive |
| **GL-0007** | 0.65 ± 0.07 | <0.0001 | Synergism |
| **GL-0008** | 1.30 ± 0.61 | 0.1052 | Additive |
| **GL-0009** | 28.69 ± 40.84 | 0.0203 | Very strong antagonism |

Combination index (CI) were computed according to Chou & Talalay method at 1.5 $EC_{50}$. Interpretation of CI is as follows: CI>10: very strong antagonism; CI 3.3-10: strong antagonism; CI 1.45-3.3: antagonism; CI 1.15-1.45 moderate antagonism; CI 0.85-1.15: additive; CI 0.85-0.65: moderate synergism; CI 0.3-0.65: synergism; CI 0.1-0.3 strong synergism; CI <0.1: very strong synergism. Two-tailed t-test have been used to assess whether CI were significantly different as compared with additivity (1.0 ± 0.15).

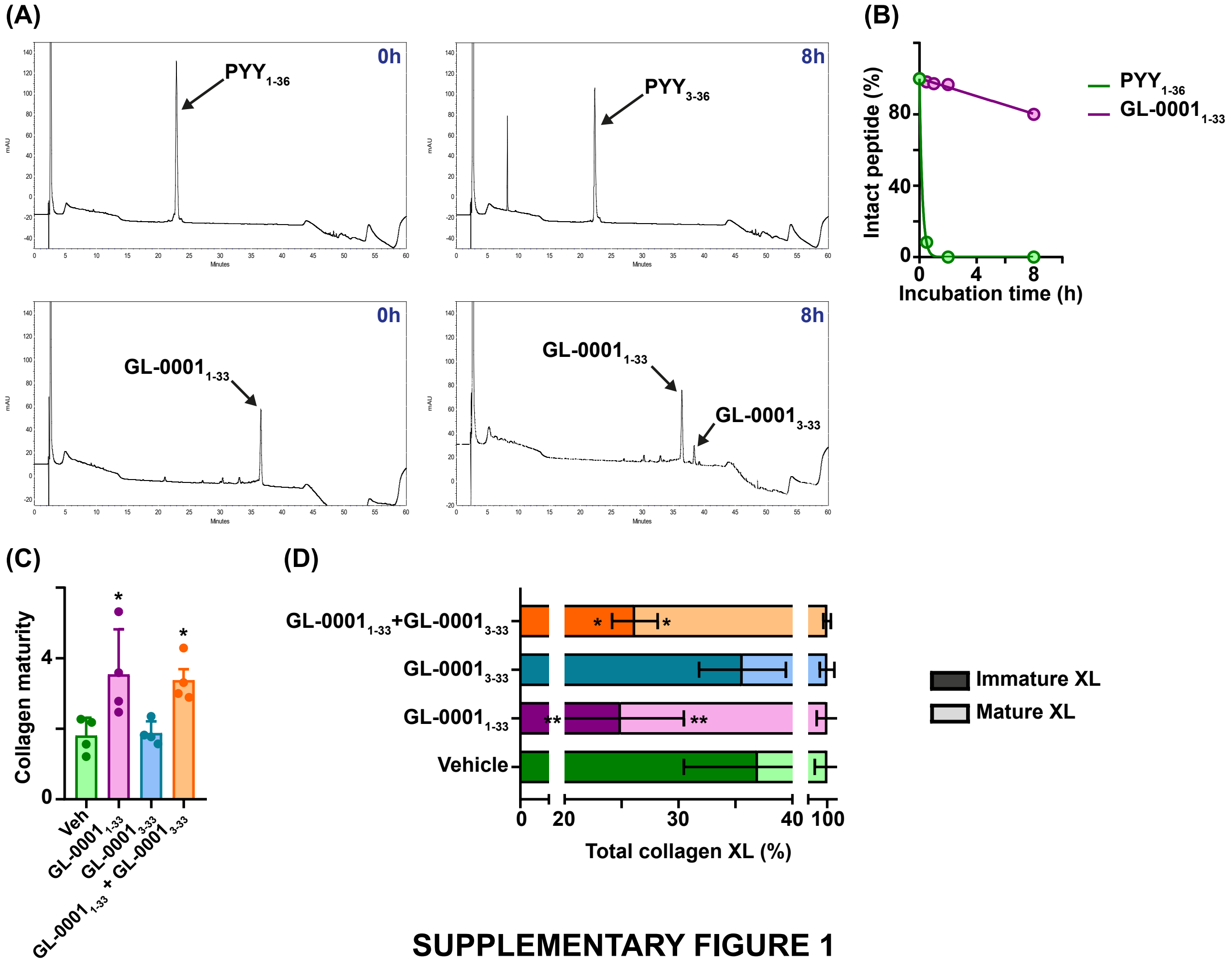


**SUPPLEMENTARY FIGURE 1**

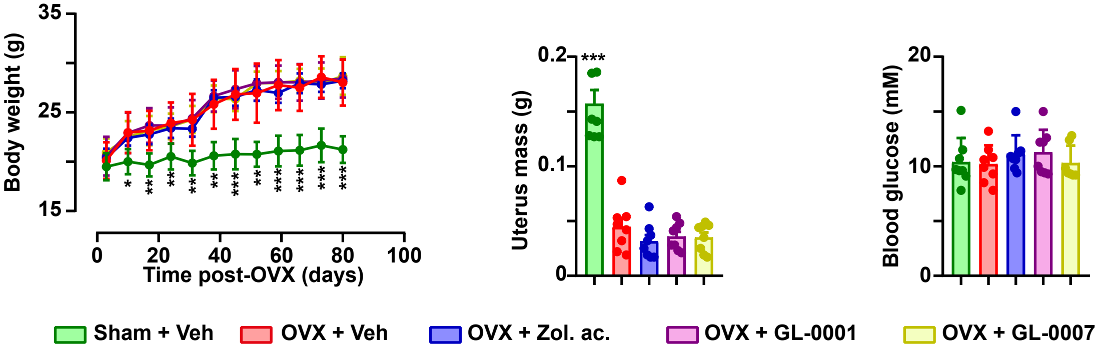


### SUPPLEMENTARY FIGURE 2

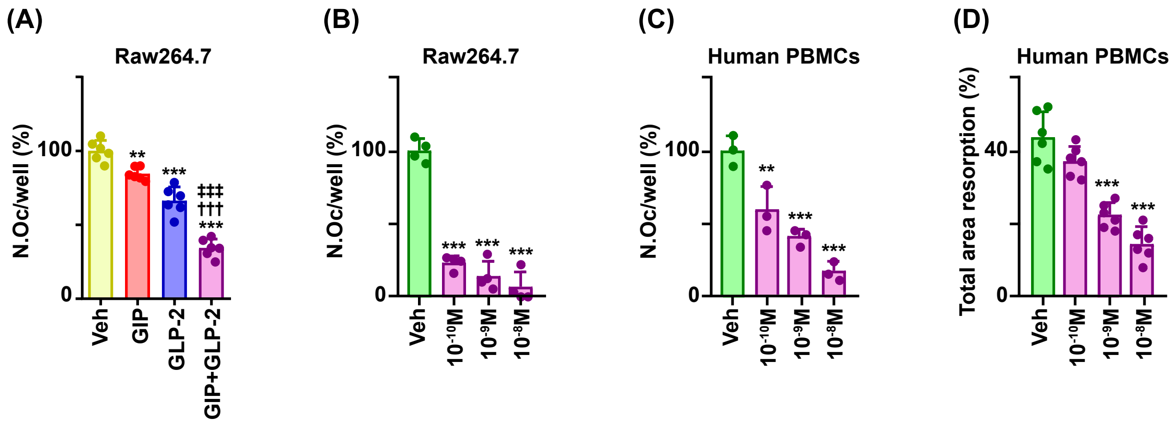


#### SUPPLEMENTARY FIGURE 3
